# Circular RNA AEBP2: A novel regulator in dendritic cell development

**DOI:** 10.1101/2024.10.23.619919

**Authors:** Qinfeng Zhou, Bowen Wang, Shuailong Li, Adam Greasley, Weiping Min, Samen Maleki Vareki, Xiufen Zheng

## Abstract

Dendritic cells (DCs) are the main antigen-presenting cells and play a central role in activating adaptive immunity. How DCs is regulated during their development remains to be further elucidated. Circular RNA (circRNA) is a new type of non-coding RNA playing a critical in various cell events. However, little is known about the role of circRNA in DCs. Using *in situ* RNA hybridization, we found that DCs expressed exonic circular RNA AEBP2 (circAEBP2). circAEBP2 was upregulated in mature DCs compared to immature DCs. Knockdown of circAEBP2 in DCs using small interference RNA (siRNA) arrested DCs at an immature state with lower expression of MHC class II, CD40, CD80, and CD83. circAEBP2 knockdown in DCs also decreased the phosphorylation of nuclear factor-kappa B p65 (P-p65) but increased CD200R1. CircAEBP2-silenced DCs had a lower capacity to activate allogeneic CD4^+^ and CD8^+^ T cells *in vitro* and induced differentiation of regulatory T cells (Tregs). RNA immunoprecipitation assays revealed that circAEBP2 interacts with heterogeneous nuclear ribonucleoprotein F (hnRNP F) which is highly expressed in DCs and subsequently prevented hnRNP F nuclear translocation. In conclusion, we demonstrate that circAEBP2 regulates the development and function of DCs and knockdown of circAEBP2 induces tolerogenic DCs (Tol-DCs) through interacting with the hnRNP F/CD200R1/P-p65 signaling pathway. circAEBP2 is a new regulator in DCs and modulating it provides a novel strategy for inducing Tol-DCs.

## Introduction

Dendritic cells (DCs) are key hematopoietic cells in regulating innate and adaptive immune responses(1-3). DCs originate from bone marrow progenitors, differentiate into immature DCs (imDCs) with low expression of major histocompatibility complex (MHC) and costimulatory molecules, and further develop into mature DCs (mDCs) in response to a stimulus(4). imDCs have a superior capacity of antigen phagocytosis but are unable to activate T cells. In contrast, mDCs highly expressing MHC class I/II and costimulatory molecules present antigenic peptides to T cells and initiate the MHC class I-restricted cytotoxic leukocyte (CTL) response and the MHC class II restrictive CD4^+^ response. DCs are the most effective professional antigen-presenting cells driving T cell activation, differentiation, and function(5-9). According to their immune function, DCs are divided into two major categories: immune-reactive and immune-suppressive. The former activates T cells and promotes T helper-1 (Th1) differentiation, thus initiating immune response, whereas the latter is unable to activate naïve T cells and enhances regulatory T cells (Tregs)(10). Generally, imDCs are immunosuppressive while mDCs are immunoreactive(11, 12). Induction of imDCs has been used to suppress immunity and to treat autoimmune disease and immune rejection in organ transplantation(11).

Multiple pathways and gene regulators including non-coding RNAs have been reported to participate in the development of DCs, determining the fate and function of DCs(13-18). Emerging evidence has shown that circular RNA (circRNA), a new type of non-coding RNA previously thought to be a byproduct during transcription, is crucial in multiple physiological and pathological processes including cell development, proliferation, and disease development by functioning as an important gene regulator (19). We have shown previously that circRNA is also involved in DC development(20). However, how circRNA modulates DCs and whether it is a novel gene regulator for DCs remains unknown.

In the present study, we investigated the role of circular RNA AEBP2 which is back-spliced from the AE Binding Protein 2 (AEBP2) gene RNA in DC development. We showed that circAEBP2 is essential for DC maturation and ability to stimulate allogeneic T cell responses. Mechanistically, we showed that circAEBP2 interacts with heterogeneous nuclear ribonucleoprotein F (hnRNP F) and subsequently prevented hnRNP F nuclear translocation in DCs, thereby regulating the development and function of DCs.

## Materials and Methods

### Animals

Male 8-10 weeks-old C57BL/6 and BALB/c mice were purchased from Charles River Laboratory (Senneville, Quebec, Canada). Animal usage was approved by the Animal Use Committee of Western University and experimental procedures were complied with the guidance of the committee.

### Dendritic cell culture

Bone marrow (BM) cells were flushed from the aseptic femur, and tibia from male C57BL/6 mice with RPMI 1640 (ThermoFisher Scientific, Mississauga, Ontario, Canada) and centrifuged for 5 min at 2000 rpm and 4°C. Collected cell pellets were lysed with 1 ml ACK lysis buffer (Sigma, Canada) to lysis red blood cells for 1 min at room temperature, followed by washing with 10 ml PBS twice. Isolated BM progenitor cells (2×10^6^ cells/well) were cultured in 6-well plates (Sarstedt, Canada) with 4 ml DC complete medium (DC-CM) consisting of RPMI 1640, 10% FBS (ThermoFisher Scientific, Canada), 100U/ml penicillin and streptomycin (Sigma, Canada), 2 mM Glutamine (Sigma, Canada), 10 ng/ml interleukin 4 (IL-4) (PeproTech, Rocky Hill, NJ) and10 ng/ml granulocyte/macrophage colony stimulating factor (GM-CSF, PeproTech) at 37°C, 5% humidified CO_2_. Half the volume of DC-CM was replaced every two days.

### Transfection with siRNA

Two siRNA sequences were designed to target the circAEBP2 junction sequence: CircAEBP2 siRNA # 1 and CircAEBP2 siRNA #2. siRNAs were synthesized by Sigma (Sigma-Aldrich Canada Co, Oakville, Ontario). Negative control (NC) siRNA was purchased from Sigma and used as a scrambled siRNA control.

DCs cultured for 4 days were collected and re-plated in a 6-well plate (2×10^6^ cells/well) with 2 ml DC medium without antibiotics one day prior to transfection. DCs were transfected with 2 µg circA EBP2 siRNA #1, circAEBP2 siRNA #2, or NC siRNA with 4 µl Endofectin (Genecopoeia, Rockville, MD) at 700 µl DC-CM without antibiotics according to the manufacturer’s instruction. Six hours after transfection, fresh DC medium without antibiotics was added to a final volume of 2 ml per well and cells were continued to culture at 37°C, 5% humidified CO_2_ for 48 hours. Another 2 ml fresh DC-CM were added to the cells at the first 24 hours after transfection.

### RNA isolation and quantitative reverse transcriptase polymerase chain reaction (qRT-PCR)

Total RNA was extracted with Trizol (Invitrogen, Mississauga, Ontario, Canada). cDNA was synthesized with random hexamer (Invitrogen, Ontario,Canada) and M-MuLV Reverse Transcriptase (New England Biolabs, Ontario, Canada) as described elsewhere(20).

Real time quantitative PCR was conducted using SYBR Green Super Mix (Bioline, Ontario, Canada) according to the manufacturer’s instruction. The PCR was performed at 95°C for 2 min, followed by 40 cycles at 95°C for 5 sec, 60°C for 10 sec, 72°C for 8 sec with CFX Connect™ Real-Time System instrument (Bio-Rad, Mississauga, Ontario, Canada). Primer sequences are listed in Supplementary Table 1. The relative expression of circAEBP2 and AEBP2 were analyzed as ΔΔCt where GAPDH is used as an internal loading control.

### DC phenotype detection

Collected DCs (2 × 10^5^) were resuspended in PBS with 2% FBS and stained with FITC-conjugated anti-CD11c mAb (eBioscience, San Diego, CA), PE-Cy5-conjugated anti-CD40 mAb (Biolegend, San Diego, CA), PE-conjugated anti-CD80 mAb (Biolegend), APC-conjugated anti-CD83 mAb (BD Biosciences, San Jose, CA), eFluor®450-conjugated anti-MHC II mAb (eBioscience) for 30 min at 4 °C or 15 min at room temperature in the dark. Stained cells were washed with PBS with 2% FBS (2% FBS-PBS) once. The fluorescence intensities of Abs bounded to DCs were measured by Cytoflex S flow cytometer (Beckman Coulter, Miami, FL).

### T cell proliferation

T cell proliferation was measured using the carboxyfluorescein succinimidyl ester (CFSE) dilution assay. T cells were isolated from splenocytes of BALB/c mice using a MojoSort mouse CD3 T cell isolation kit (Biolegend) according to the manufacturer’s instructions. Purified T cells were then labeled with 5 µM CFSE (BD Biosciences) in PBS for 15 min at 37°C, followed by washing with PBS once. CFSE-labeled T cells (2 ×10^5^/well) were co-cultured with day 7 allogeneic DCs (40,000/well) from C57BL/6 mice at a ratio of T: DC= 5: 1 in a 96 well round bottom plate for 96 hours at 37°C, 5% humidified CO_2_. T cells were then stained with PE-conjugated anti-CD4 mAb (Biolegend) and PE-Cy5-conjugated anti-CD8 mAb (Biolegend) for 30 min at 4°C prior to flow cytometry analysis. The fluorescence intensity of CFSE and Abs bounded to T cells were measured by Cytoflex S flow cytometer.

### CD4^+^CD25^+^Foxp3^+^ Treg generation and detection

T cells isolated from splenocytes of BALB/c mice using a MojoSort mouse CD3 T cell isolation kit (2 ×10^6^/well) were co-cultured with day 7 DCs (2 × 10^5^/well) with 2 ml RPMI 1640 medium with 10% FBS and antibiotics in a 24-well plate for 5 days at 37°C, 5% humidified CO_2_. Cells were collected and stained with PE-Cy5-conjugated anti-CD4 mAb (Biolegend), PE-Cy7-conjugated anti-CD25 mAb (Biolegend) for 15 min at room temperature in the dark, followed by washing with PBS buffer with 2% FBS once. Cells were then fixed and permeabilized using a fix/permeabilization kit (ThermoFisher Scientific, Mississauga, Ontario, Canada) according to the manufacturer’s instructions. Permeabilized cells were further stained with an Alexa Fluor® 488 Foxp3 antibody (Biolegend) for 30 min at 4°C in the dark, followed by washing with 2% FBS-PBS. Cells were suspended in 2% FBS-PBS and subjected to flow cytometry analysis using a Cytoflex S flow cytometer.

### Western blotting

Cells were lysed using radioimmunoprecipitation assay (RIPA) buffer (Cat. # 9806S, Cell Signaling Technology (CST), Danvers, MA) supplemented with protease inhibitor phenylmethylsulfonyl fluoride (PMSF, cat. # 8553S, CST). Protein concentrations were measured using Quick Start™ Bradford 1x Dye Reagent (Bio-Rad). Proteins (25 µg) were separated by 12% polyacrylamide gel electrophoresis (PAGE, Sigma), and electro transferred onto polyvinylidene fluoride (PVDF) membranes (Bio-Rad) using Trans-Blot® Turbo™ Transfer System (Bio-Rad) according to the manufacture’s instruction. The membranes were blocked with TBS buffer containing 5% milk for 1 hour at room temperature, then blotted with AEBP2 (D7C6X) mAb (1:1000, Cat. # 14129S, Cell Signaling), P-NFκB P65 mAb (1:2000, Cat. # 3033L, CST), β -Actin mAb (1:4000, Cat. # G0512, Santa Cruz Biotechnology (SCB), San Diego, CA), hnRNP F mAb (1:800, Cat. # ab50982, Abcam) overnight at 4°C. Blotted membranes were washed with 0.5% Tween TBS for 10 min, 3 times, followed by blotting with goat anti-mouse (1:4000, Cat. # sc-2005, SCB), or donkey anti-rabbit horseradish peroxidase conjugated secondary antibody (1:4000, Cat. # sc-2313, Snata Cruz). Proteins were developed with Clarity™ Western ECL Substrate (Cat. # 170-5060, Bio-Rad) or Clarity™ Max Western ECL Substrate (Cat. # 170-5062S, Bio-Rad) by FluorChem M system (Proteinsimple, Santa Clara, CA) following manufacturer’s protocol. Protein band intensities were measured with ImageJ software(21).

### Immunocytochemistry (ICC)

DCs (20,000/well) were plated in a 24-well plate pre-coated with 1% gelatin (Sigma). Eighteen hours after culture, cells were washed with cold PBS twice, and fixed with 500 µl 4% paraformaldehyde (PFA, Sigma) for 20 min at RT. 200 µl 0.1% Triton (Sigma) PBS were added to the cells for 15 min at room temperature. AEBP2 (D7C6X) rabbit mAb (1:50 dilution), P-NFκB P65 rabbit mAb (1:50), and hnRNP F rabbit mAb (1:100) were incubated for 1 hour at room temperature. After washing with PBS for three times, cells were incubated with Alexa Flour 488 goat anti-rabbit IgG (H+L) (Cat. # A11008, ThermoFisher Scientific) in the dark for 1 hour at room temperature. Cells were then counterstained for 5 min in the dark with 1:10,000 diluted 4, 6-diamidino-2-phenylindole dihydrochloride (DAPI, ThermoFisher Scientific) and imaged under a fluorescence microscope (x 200).

### Synthesis of biotinylated probes

Biotinylated probes were synthesized by PCR using Biotin-14-dCTP. Briefly, the synthesis of biotin labeled probes was conducted by PCR in a 50 µl reaction volume with one Taq® standard reaction buffer (Biolabs), 0.2 mM dNTPs where dCTP: Biotin-14-dCTP =1:1, 8nM DNA oligo template probes, 0.2 µM forward/reverse primers, 10U DNA Taq DNA polymerase recombinant (ThermoFisher Scientific). PCR was performed at 95°C for 3 min, followed by 50 cycles at 95°C for 5 sec, 46°C for 10 sec, 72°C for 10 sec, with a final step at 72°C for 10 min in a T100™ Thermal Cycler instrument (Bio-Rad). DNA oligo template sequences for circAEBP2 probe and random probe and PCR primer sequences are listed in Supplementary Table 1.

### RNA immunoprecipitation assay (RIP)

DCs (1×10^7^) were collected, washed with cold PBS once, and then lysed with 200 µl 1x RIPA buffer containing protein inhibitor PMSF on ice for 20 min and vortexed twice during lysis. Lysed cells were centrifuged at 14,000 rpm for 20 min at 4°C. Supernatant was transferred to a new 1.5 ml Eppendorf tube on ice. 3 µg biotinylated denatured DNA oligo probes were denatured for 5 min at 95°C, chilled 5 min on ice, and added into the cell lysate and incubated for 2 hours at room temperature with gentle agitation. 50 µl pre-washed streptavidin C1 Dynabeads (ThermoFisher Scientific) were added to the above cell lysate-DNA probe mixture and incubated for 1 hour at room temperature, followed by washing with 100 µl co-IP buffer (ThermoFisher Scientific) 5 times. 30 µl RNase-free H_2_O was added to the tube and incubated at 65°C for 10 min to elute DNA probe bound compounds. The eluted compounds were subjected to RNA and protein isolation using QIAzol (Qiagen, Misusage, ON, Canada) following the manufacturer’s instruction. Briefly, chloroform (1/5 volume of QIAzol solution) was added onto QIAzol solution, mixed, and centrifuged at 14,000 rpm for 15 min. The top aqueous layer was collected to extract RNA using isopropanol precipitation. To extract protein from the bottom organic layer, 0.3 ml of 100% ethanol was added per ml of QIAzol and mixed inversely, then left at room temperature for 3 min. After centrifugation at 14,000 rpm for 5 min at 4°C, the top layer was transferred to a new microtube. 2x volume of isopropanol was added to the tube, mixed, incubated for 10 min at room temperature, and then centrifuged at 12,000g for 10 min at 4°C. The protein pellet was washed with at least 500 µl 95% ethanol twice, air dried, and dissolved with 1% SDS solution for 1 hour at 50°C. Samples were ready for mass spectroscopic analysis or western blotting.

### Immunoprecipitation assay (IP)

Cells were lysed using RIPA buffer (CST) supplemented with protease inhibitor PMSF (CST). Protein concentrations were measured by using Quick Start™ Bradford 1x Dye Reagent (Bio-Rad). 600 µg total proteins were used for IP. 5 µg hnRNP F specific Abs or IgG which is unable to bind to hnRNP F as a control were added into the above cell lysate and incubated for 4 hours at 4°C with gentle agitation. 20 µl Protein A/G Plus-Agarose (Cat. # SC-2003, Santa Cruz) were added to the mixture and incubated overnight at 4°C. Samples were centrifuged at 10,000 rpm for 10 min at 4°C, and then washed with PBS for three times and centrifuged at 10,000 rpm for 10 min. 150 µl QIAzol regent was added to the IP compound to isolate RNA and protein, followed by qRT-PCR and Western blotting for detection of RNA and protein, respectively.

### Mass spectrometry (MS)

Protein samples were reduced with DTT, alkylated with iodoacetamide, and digested overnight with trypsin. Digested samples were cleaned up using a homemade C18 spin column. Cleaned samples were lyophilized. Samples were then resuspended in 12 µl buffer A (0.1% FA). 6 µl of each sample was analyzed by nanoflow liquid chromatography on an Ultimate 3000 LC system (ThermoFisher Scientific) online coupled to a Fusion Lumos Tribrid mass spectrometer (ThermoFisher Scientific) through a nano electrospray flex-ion source (ThermoFisher Scientific). Samples were loaded onto a 5 mm µ-precolumn (ThermoFisher Scientific) with 300 µm inner diameter filled with 5µm C18 PepMap100 beads. Peptides were then subjected to chromatography on a 15 cm column with 75 µm inner diameter with 2 µm reverse-phase silica beads. Peptides were separated and directly electrosprayed into the mass spectrometer using a linear gradient from 4% to 30% ACN in 0.1% formic acid over 160 min at a constant flow of 300 nl/min. The linear gradient was followed by a washout with up to 95% ACN to clean the column followed by an equilibration stage to prepare the column for the next run. The Fusion Lumos was operated in data-dependent mode, switching automatically between one fill scan and subsequent MS/MS scans of the most abundance peaks with a cycle time of 3 seconds. Full Scan MS1s were acquired in the Orbitrap analyzer with a resolution of 120,000, scan range of 400-1,600 m/z. The maximum injection time was set to 50 milliseconds with an AGC target of 4e5. RF lens was set to 30. The fragment ion scan measured in the linear ion trap using a Quadrupole isolation window of 0.7 m/z, and HCD fragmentation energy of 30. Ion trap scan rate was set to turbo, with a maximum ion injection time of 22 milliseconds and an AGC target set to 1e4.

### *In situ* RNA Hybridization

Cultured DCs (2×10^5^/well) were plated in a 24-well plate which was pre-coated with 1% gelatin for at least 2 hours at 37°C and cultured overnight. Cell were washed with cold PBS twice and fixed with 500 µl 4% PFA for 20 min at room temperature. 200 µl of 0.1% Triton (Sigma) diluted in PBS were added to permeabilize cells for 15 min at room temperature. Denatured RNA-biotinylated probes (1.5 μg) were added to preheated 150 µl hybridization buffer (containing 2% methanamide, 0.5 M NaCl, 0.1 M Tris-HCl, 1% SDS) at 55°C. The mixture was immediately added to the fixed/permeabilized cells and incubated for 2 hours at room temperature. Cells were washed with preheated hybridization wash buffer containing 0.5mo/L EDTA at 55°C for 15 min. 1 µg/ml Alexa Fluor™ 594 conjugated streptavidin (Cat. # S11227, Invitrogen) was added to cells and incubated for 1.5 hours at room temperature in the dark. The hybridized cells were counterstained with DAPI (1:5000) for 5 min in the dark and imaged using fluorescent microscopy (x 200).

### Statistical analysis

The mean with standard deviation was used to present data. Statistical analysis of experiments was performed using GraphPad Prism (2016, Version 7.0a). Paired and unpaired Student’s t test or one-way ANOVA was applied for comparisons between two groups, or among three groups or more groups. *P* value < 0.05 was considered statistically significant.

## Results

### CircAEBP2 expression in DCs

First, we determined the expression of circAEBP2 in DCs. Bone marrow derived DCs (BM-DCs) were cultured *in vitro*. RNA was extracted on days 2, 4, 6, and 8 of culture, and the amount of circAEBP2 was determined by quantitative reverse transcriptase-polymerase chain reaction (qRT-PCR) using divergent primers. It was found that circAEBP2 levels gradually increased in DCs over culture time and circAEBP2 was expressed significantly greater in DCs collected on day 8 than those collected earlier (approximately 3 fold greater than on day 2)(Figure 1A).

**Figure 1.**
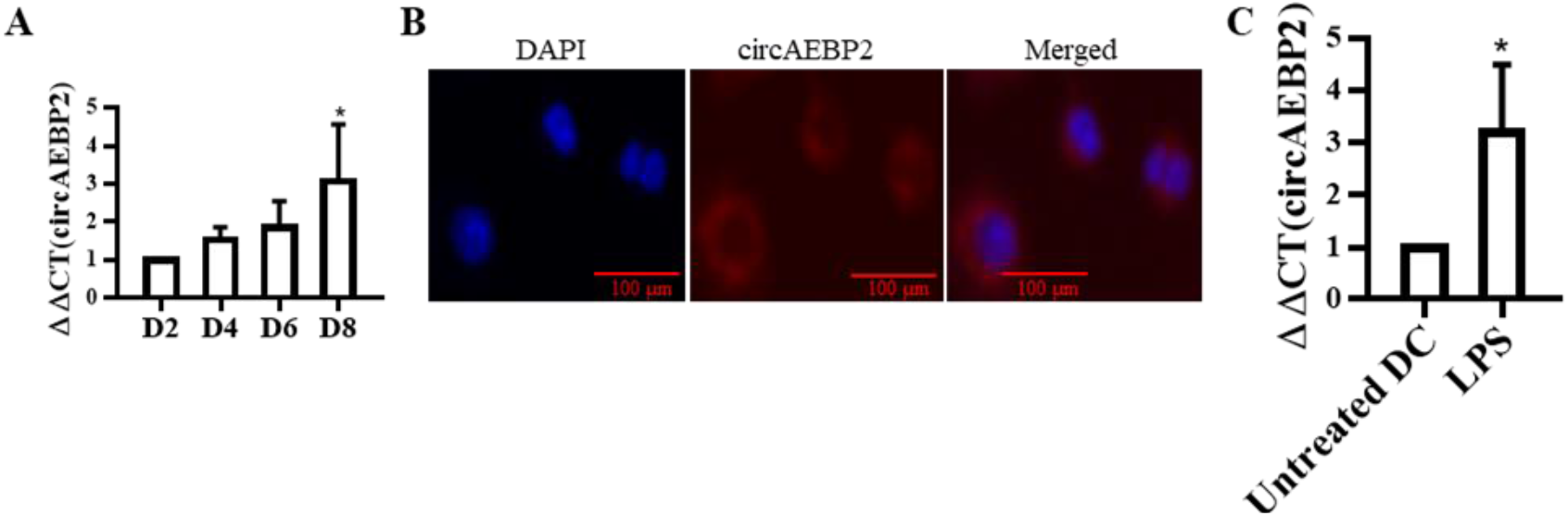
circAEBP2 and AEBP2 expression in DCs. (A) circAEBP2 expression in DCs. BM-derived DCs from C57BL/6 were cultured in DC-medium in *vitro*. RNA was collected from DCs on the indicated days. The relative expression of circAEBP2 were measured by qRT-PCR. n=4, * *p* < 0.05 vs. DC day 2 (B) Distribution of circAEBP2 in DCs. *In situ* RNA hybridization was conducted using biotinylated circAEBP2 probes, followed by visualization with Alexa Fluor™ 594-streptavidin. Scale bar, 100µm. (C) LPS increased the expression of circAEBP2 in DCs. On day 7 of culture, DCs were stimulated with 100 ng/ml LPS for 1 hour. circAEBP2 expression was measured by qRT-PCR. n=3, * p < 0.05 vs untreated DC.

Next, to determine distribution of circAEBP2 in the cells, *in situ* RNA hybridization was conducted using biotinylated circAEBP2 probes which were prepared by PCR with biotin-14-dCTP and complementary to the junction sequence of circAEBP2, followed by visualization with Alexa Fluor™ 594-streptavidin. We found that circAEBP2 was exclusively located in the cytoplasm (Figure 1B), not in the nucleus.

Furthermore, we investigated the effect of lipid polysaccharide (LPS) stimulation on the expression of circAEBP2. To do this, BM-DCs were treated with 100 ng/ml LPS for 1 hour on day 7 and then RNA was extracted. LPS stimulation increased the level of circAEBP2 in DCs compared to untreated day 7 DCs approximately 3-fold as measured by qRT-PCR (Figure 1C).

### Knocked down of circAEBP2 reduces the maturation of DCs

To determine the effect of circAEBP2 on DC development, DCs maintained in culture for 5 days were transfected with two different small interference RNAs (siRNA, #1 and #2) specifically targeting the circAEBP2 RNA conjunction sequence or control negative siRNA, respectively. The result of qRT-PCR showed that the level of circAEBP2 RNA 48 hours post-transfection was reduced to 50 % in these DCs transfected with either circAEBP2 siRNA #1 or #2 compared with negative control (NC) siRNA (Figure 2A).

**Figure 2.**
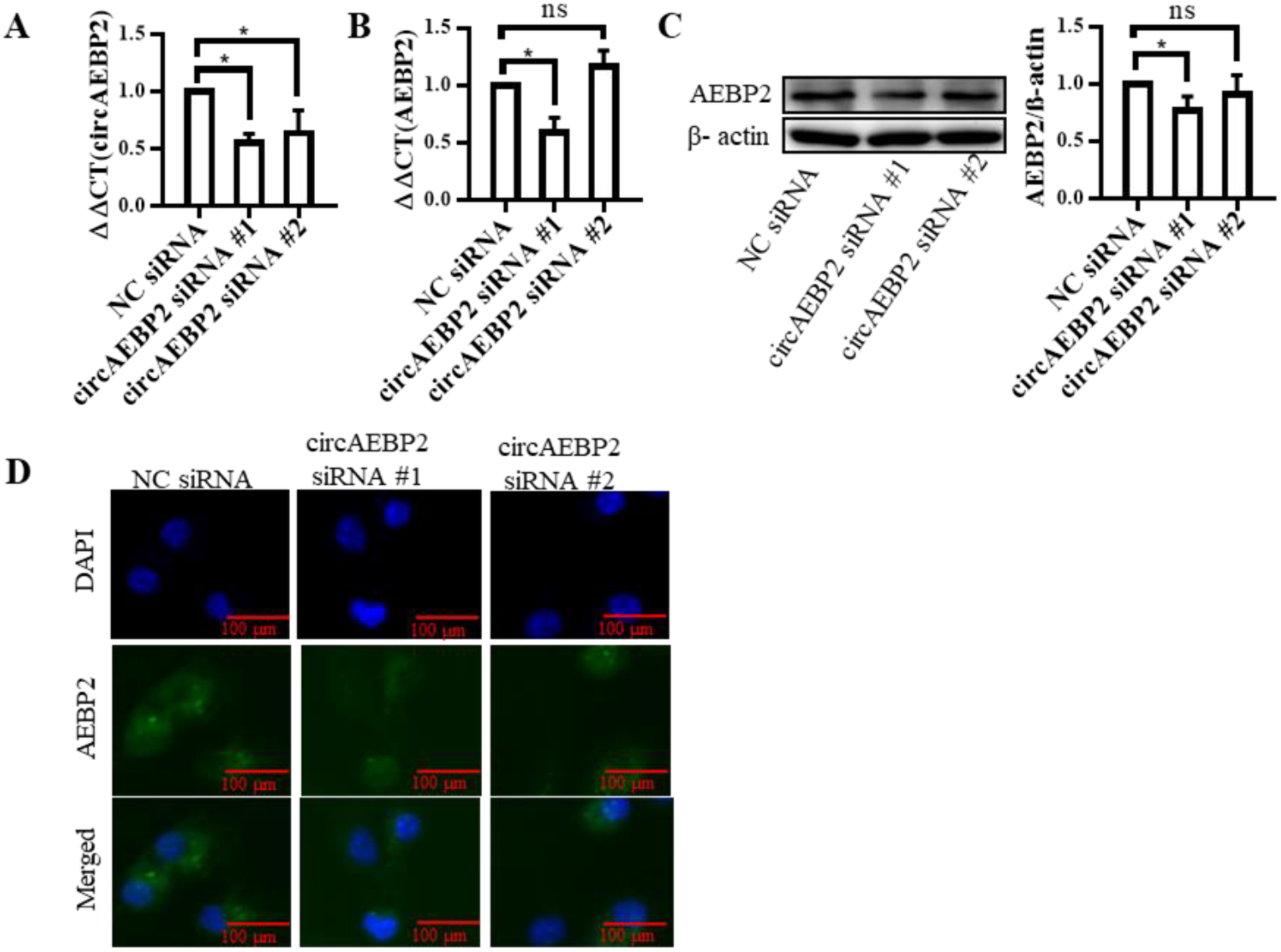
Knocking down circAEBP2 and AEBP2 in DCs with siRNA. (A) siRNA downregulation of circAEBP2 in BM-DCs. On day 5, cultured DCs were transfected with circAEBP2 siRNA #1, or circAEBP2 siRNA #2, or negative control siRNA for 48 horns. The relative expression of circAEBP2 were measured by qRT-PCR and normalized with NC controls. (B) The effects of siRNAs on AEBP2 expression. n=4, * *p* < 0.05, ns *p* > 0.05. (C) AEBP2 protein expression. Western blotting was conducted to determine AEBP2 protein expression using AEBP2 Abs. Left panel: representative images; right panel: semi-quantitative results of western blotting, β-actin was used as a loading control, the ratio of AEBP2/actin were calculated and normalized with NC controls. n=6, \**p*< 0.05, ns *p*> 0.05. (D) Immunofluorescent staining images of AEBP2. Day 7 DCs were cultured in a 1% gelatin pre-coated plate. Cells were fixed, permeabilized and blotted with AEBP2 Abs and Alexa Flour 488 labelled Abs. Images were taken under a fluorescent microscope at the magnification of x 200. Green: AEBP2. Blue: DAPI. Scale bar. 100µm.

We also observed that the level of AEBP mRNA was downregulated by approximately 60 % in DCs transfected with circAEBP2 siRNA #1 (Figure 2B). However, transfection with circAEBP2 siRNA #2 did not change the levels of AEBP2 mRNA (Figure 2B) in DCs compared with NC siRNA (Figure 2B). Western blotting confirmed the effects of the siRNAs on AEBP2 expression at the protein level (Figure 2C). An immunocytochemistry (ICC) assay with AEBP2 Abs was also conducted to measure AEBP2 protein. The ICC results were consistent with western blotting results (Figure 2D). Therefore, we selected circAEBP2 siRNA #2 which specifically knocked down circAEBP2 sparing AEBP2 mRNA for forward experiments and referred it as circAEBP2 siRNA.

Next, we determined the effect of circAEBP2 on DC maturation by transfecting DCs with circAEBP2 siRNA and NC siRNA for 48 hours. We then examined DC maturation markers using flow cytometry. The expression of MHC II, costimulatory molecules CD40, CD80 and CD83 (Figure 3A-D), was significantly decreased in DCs transfected with circAEBP2 siRNA compared to NC siRNA.

**Figure 3.**
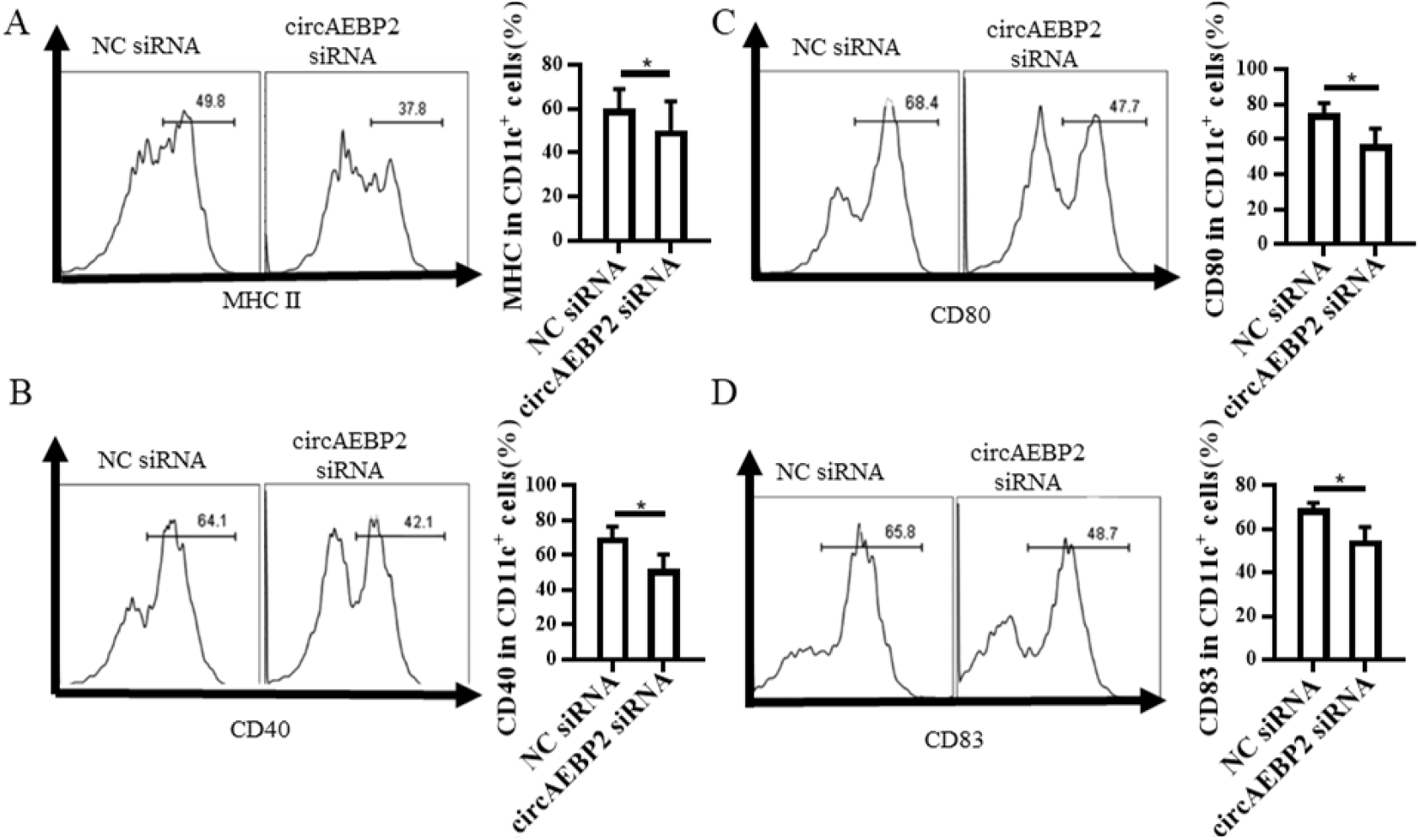
Knockdown of circAEBP2 reduces the maturation of DCs. DCs were transfected with 2μ circAEBP2, or NC siRNA on day 5 using endofectin for 48 hours. DCs were stained with MHC II eFluor®450 Abs, CD11c-FITC Abs, CD40 PE-Cy5 Abs, CD80 PE Abs, CD83 APC Abs. followed by flow cytometry analysis. Viable CD11c positive cells were gated. MHC II (A), CD40 (B), CD80 (C), and CD83 (D). Left panel: representative original flow cytometric histogram graphs. Right panel: Summarized results of flow cytometry from n = 5 experiments, * *p* < 0.05.

### CircAEBP2 regulates the NFκB pathway and CD200R1

The nuclear factor κB (NFκB) pathway is essential for DC maturation and upregulation of DC costimulatory molecules such as CD80 and CD40 requires NFκB activation(22). Accordingly, we determined the effect of circAEBP2 downregulation on the expression of phosphorylated p65 (P-p65) or Rel A which is a member of the NFκB pathway by western blotting (Figure 4A) and ICC (Figure 4B). We found that silencing circAEBP2 reduced the level of P-p65 by approximately 55% compared to treatment with NC siRNA. The ICC results further confirmed that circAEBP2 siRNA significantly decreased the expression of P-p65 both in the cytoplasm and nucleus of transfected cells compared with NC siRNA.

**Figure 4.**
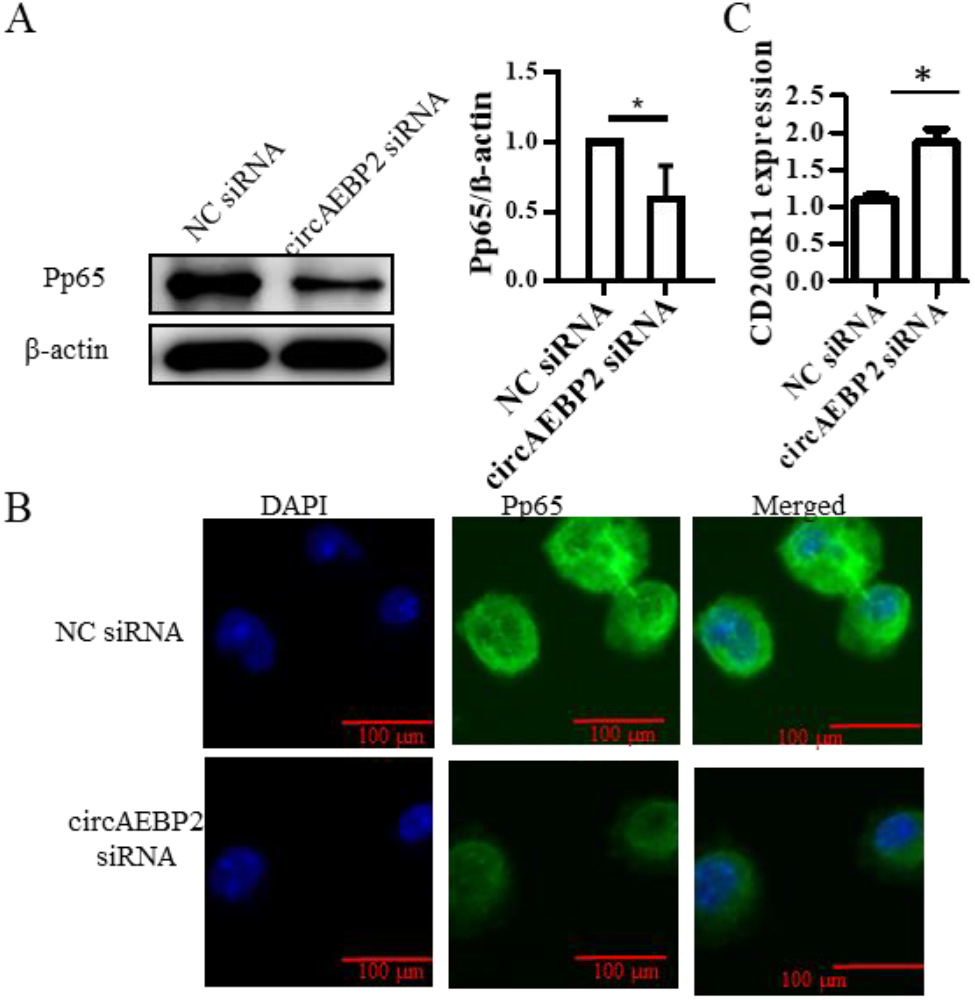
Knockdown of circAEBP2 inactivates the NFκB pathway and increases CD200R1. (A) P-p65 expression measured by Western blotting. DCs were transfected with siRNA as described above. After transfection for 48 hours, P-p65 expression was detected by Western blotting. Left panel: representative images of western blotting; right panel: semi-quantitative results of western blotting, β-Actin was used as a loading control, the ratio of Pp65/actin were calculated and normalized with NC controls. n=6, * *p* < 0.05. (B) Immunocytochemistry staining images of Pp65. Transfected DCs were plated in a 1% gelatin coated plate and subjected to ICC. Representative iminunofluorescent staining images. Green: Pp65, blue: DAPI. Scale bar, 100 µm. (C) CD200R1 expression.

Additionally, mass spectrometry (MS) analysis was conducted to determine the protein level in DCs. It was found that 40 proteins including CD200R1 which is a receptor of CD200 are increased in circAEBP2 silenced DCs (Figure 4C) compared with the control siRNA transfected DCs (*p* value < 0.05) (Supplemental Table 2).

### CircAEBP2 siRNA disables DCs to activate naïve allogeneic T cells but enhances T_reg_ generation

To determine the effect of circAEBP2 on the capacity of DCs to activate naïve T cells, a mixed lymphocyte reaction (MLR) was conducted where T cell proliferation was measured by carboxyfluorescein succinimidyl ester (CFSE) dilution assays. circAEBP2 siRNA-transfected DCs were used as stimulator cells and allogeneic CFSC-labelled naïve T cells as responder cells in the MLR. As Figure 5A shows, CD4^+^ T cell proliferation exhibited by a percentage of CFSE^dim^ CD4^+^ T cells was significantly lower when circAEBP2 siRNA-transfected DCs were used as stimulator cells (39.57 ± 2.51%) than when NC siRNA-transfected DCs were used to simulate T cells (58.13 ± 4.40%). The proliferation of CD8^+^ T cells presented a similar trend to CD4^+^ T cells (Figure 5B).

**Figure 5.**
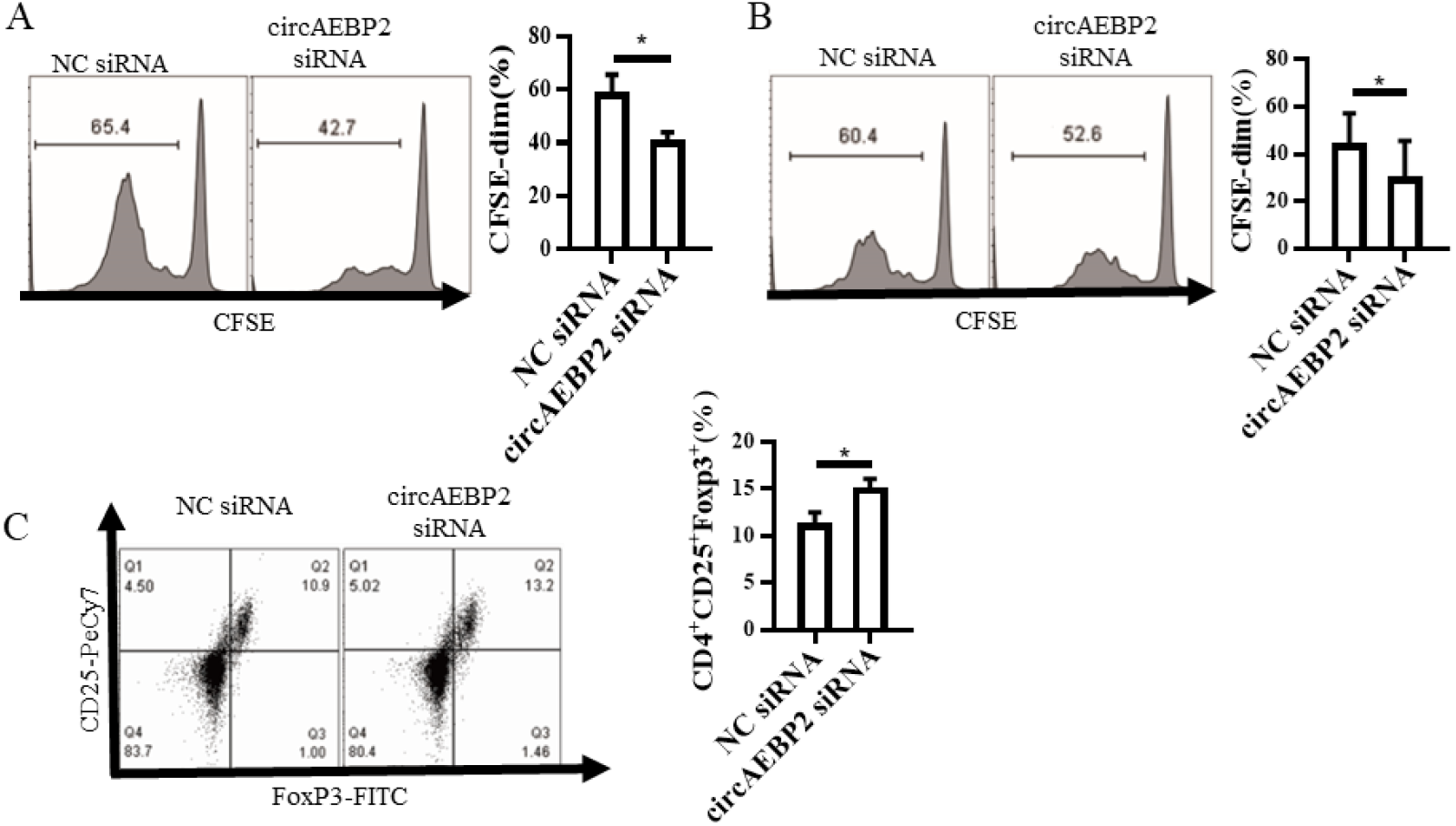
CircAEBP2 siRNA disables DCs to activate naïve allogeneic T cells and enhances Tregs generation. (A and B) T cell proliferation. siRNA transfected Day 7 DCs were co-cultured with CFSE labelled allogeneic naïve T cells at the ration of l:5=DCs: T cells for 96 horn’s. Cells were stained with CD4-PE and CD8 PE-Cy5 Abs and subjected to flow cytometry. Cells were first gated on CD4^+^ T cells (A) and CD8^+^ T cells (B). CFSE^dim^ cells represent proliferative cells while CFSE high represent non-proliferative cells. Left panel: Representative histogram graph. Right panel: summarized results of the percentage of CFSE^dim^ cells. n=4, * *p* < 0.05. (C) CD4^+^CD25^+^Foxp3^+^Tregs. siRNA transfected Day 7 DCs were co-cultured with llogeneic naïve T cells at the ratio of l:10=DCs: T cells for 5 days. Cells were stained with CD4. CD25 and Foxp3 Abs, the intensity of fluorescence was easured by flow cytometry. Cells were gated on viable CD4^+^ cells. Left panel: representative dot plot results of CD25 andFoxp3. Right panel: summarized results of flow cytometry. n=6, * *p* < 0.05.

Regulatory T cell (T_reg)_ generation is another important immune function of DCs. Accordingly, DCs transfected with siRNA for 48 hours were co-cultured with T cells for 5 days. Induced CD4^+^CD25^+^Foxp3^+^ T_regs_ were determined by flow cytometry. The results show that circAEBP2 siRNA-transfected DCs increased the percentage (∼15%) of CD4^+^CD25^+^Foxp3^+^ T_regs_ compared with NC siRNA-transfected DCs (∼10%) (Figure 5C).

### CircAEBP2 directly interacts with hnRNP F

*In situ* RNA hybridization was first performed on DCs cultured for 7 days to validate the designed biotinylated DNA probes for an RNA immunoprecipitation (RIP) assay which is used to understand how circAEBP2 functions. As shown in Figure 6A, very strong bright red fluorescence from Alexa Fluor 594 bound to biotinylated probes was observed in the DCs hybridized with circAEBP2 probes, whereas no red fluorescence were seen in the DCs hybridized with the random probes, indicating that circAEBP2 probe (and not the random probe) can bind to circAEBP2 and that the probes can be used for RIP.

**Figure 6.**
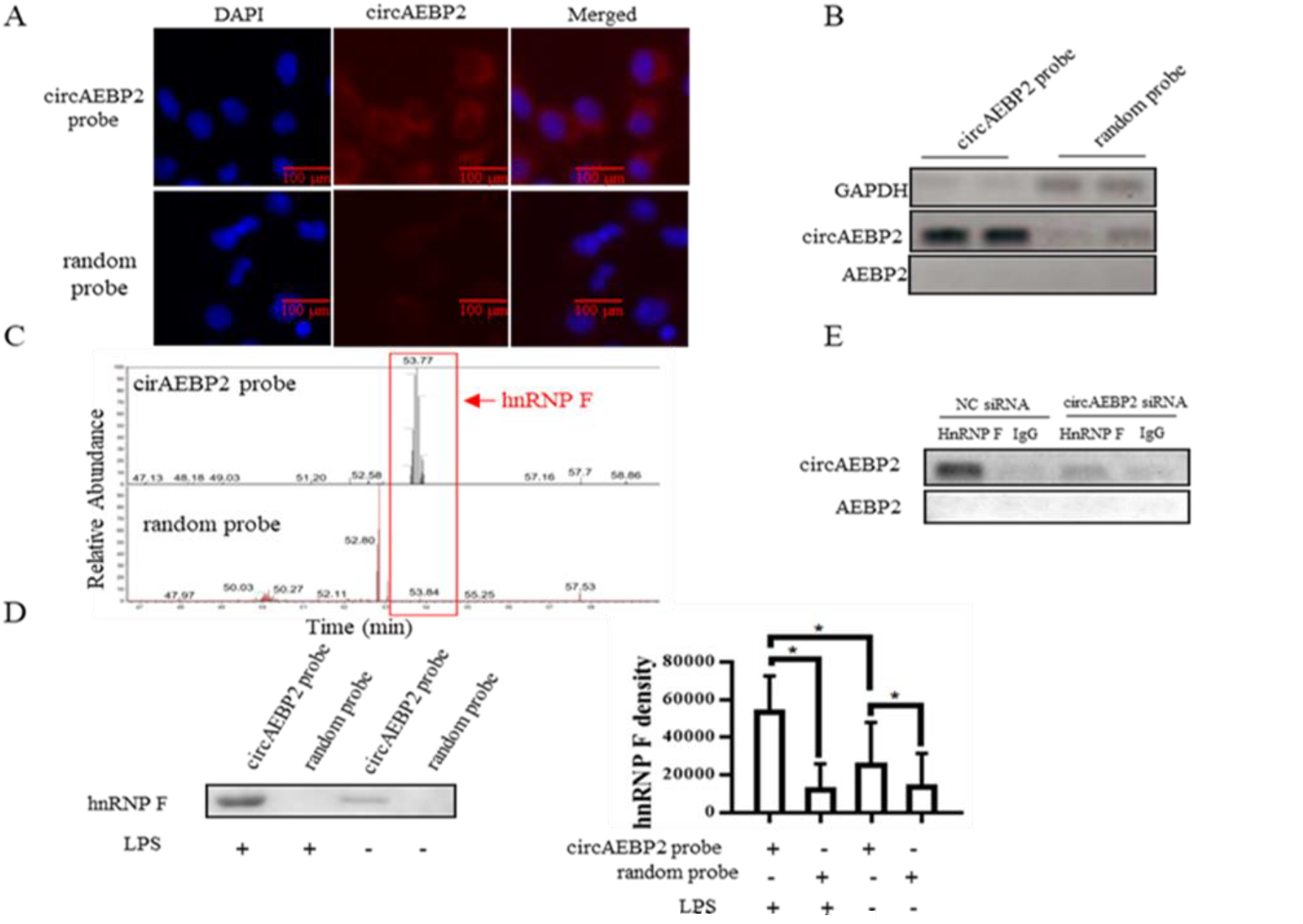
CircAEBP2 directly interacts with hnRNP F. (A) Biotinylated circAEBPZ probe not random probe detected circAEBP2. Representative inununofluorescent RNA Hybridization images of circAEBP2 in DCs. DCs were hybridized with biotinylated circAEBP2 probe or biotinylated random probe respectively, visualized with Streptavidin conjugated Alex Flour 594. Imaging was conducted by microscopy at the magnification x 200. Red: circAEBP2; Blue: DAPI for nuclei. Scale bar. 100μm. (B) CircAEBP2 pulled down by biotinylated circAEBP2 probe. 10^7^ LPS treated DCs were lysed and subjected RIP assays using biotinylated circAEBP2 probe or random probe. RNA was extracted from the Pulldown, the circAEBP2 levels were easured by RT-PCR. and PCR products were detected by electrophoresis with a 1.5% agarose gel. (C) HnRNP F unique peptide of total ion chromatography dentified in tlie cricAEBP2 probe sample. MS analysis was conducted to determine proteins in RIP pulldown. The part in tlie red box represents a peptide segment of hnRNP F. (D) CircAEBP2 interacts with hnRNP F. Proteins in RIP pulldown were extracted and subjected to Western blotting using HnRNP F Abs. Left panel: representative images: right panel: semi-quantitative results of western blotting. n=3. * p < 0.05. (E) CircAEBP2 pulled down by hnRNP F Abs. An IP assay using linRNPF Abs was performed on DC lysate. The levels of AEBP2 and circAEBP2 were detected by RT-PCR. The PCR products were detected by 1.5% agarose gel electrophoresis and visualized by EB staining.

Next, we conducted RIP assays with cell lysate from DCs stimulated by LPS using the above validated biotinylated circAEBP2 probes and random probes as LPS could increase the expression of circAEBP2 in DCs. circAEBP2 in the pulldown complex was detected by RT-PCR, followed by an agarose gel electrophoresis. The result shows that more circAEBP2 existed in the RIP complex pulled with the circAEBP2 probes than those pulled with the random probes (Figure 6B). No linear AEBP mRNA was detected in the pulldown either with the circAEBP2 probes or the random probes.

To determine which proteins were present in the RIP complex, MS was conducted. The MS results showed that hnRNP F protein was remarkably enriched by circAEBP2 probes compared with the random probes (Figure 6C). Western blotting using Abs against hnRNP F protein was also conducted to confirm the MS result. The western blotting result shows that the levels of hnRNP F protein were higher in the RIP complex treated with the circAEBP2 probes than those treated with the random probes (Figure 6D). We also found that more hnRNP F proteins were detected in the pulldown complex from the lysate of LPS-stimulated DCs than those from DCs without LPS treatment (Figure 6D).

To further confirm the interaction between circAEBP2 and hnRNP F protein, another IP assay was conducted using hnRNP F Abs and Sepharose A/G on lysate from circAEBP2 silenced DCs with circAEBP2 siRNA or control unsilenced DCs with NC siRNA. IgG protein was used as a control protein. CircAEBP2 amount in the pulldown complex was detected by RT-PCR, followed by agarose gel electrophoresis. The results show that more circAEBP2 was detected in the pulldown complex with hnRNP F Abs than those with IgG control and that less circAEBP2 was pulled down by hnRNP F Abs in the DCs silenced with circAEBP2 siRNA compared with those in NC siRNA transfected DCs (Figure 6E). Linear AEBP2 was not detected in the complex pulled down by hnRNP F Abs nor IgG proteins.

### CircAEBP2 stabilizes cytosol hnRNP F protein preventing hnRNPF nuclear translocation

To explore the distribution of hnRNP F in DCs, we conducted *in situ* immunofluorescent staining using Abs against hnRNP F and Alexa Fluro 488 labelled 2^nd^ Abs. We found that hnRNP F was expressed in both the cytoplasm and the nucleus of DCs. CircAEBP2 siRNA reduced the expression of cytosol hnRNP F compared with NC siRNA, but increased nuclear hnRNP F levels (Figure 7A). There were no obvious changes in the relative intensity of total hnRNP F (Figure 7B). However, the ratio of [nuclear hnRNP F]/ [cytosol hnRNP F] was increased (∼2.5 fold) in the circAEBP2 silenced DCs compared with the NC siRNA-transfected DCs (Figure 7C). Western blotting was also performed to measure total HnRNP F protein levels, revealing that total levels of hnRNP F protein remained unchanged between circAEBP2 siRNA-silenced DCs and NC siRNA-transfected DCs (Figure 7D), in accord with ICC data (Figure 7B).

**Figure 7.**
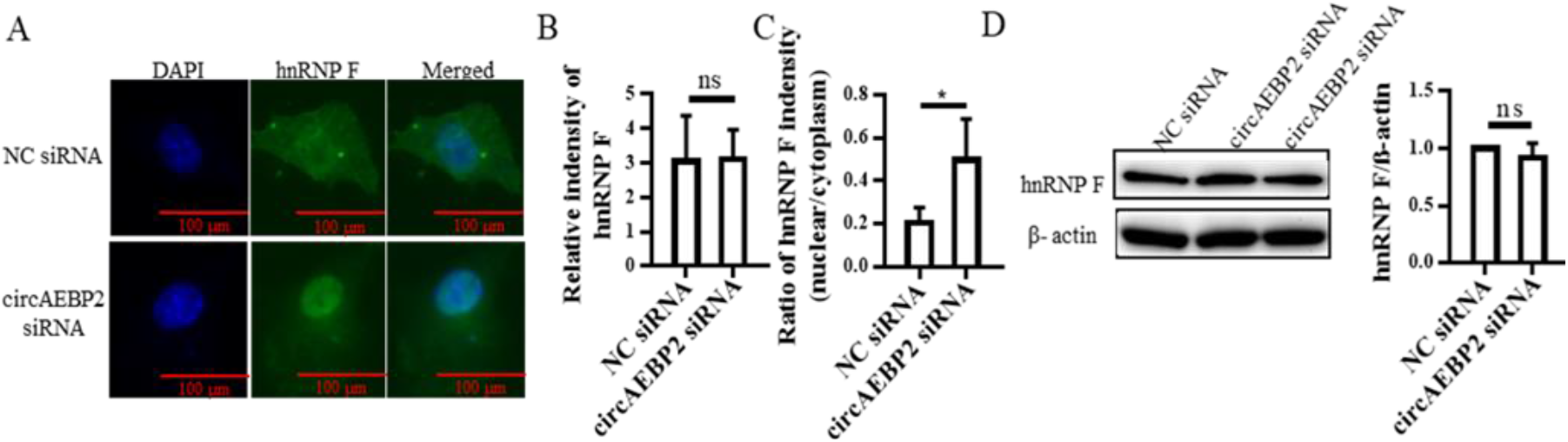
CircAEBP2 stabilizes cytosol linRNPF preventing hnRNP F nucleic translocation. (A) Immunofluorescent staining of hnRNP F. Green: linRNP F: blue: DAPI for the nucleus. Scale bar. 100μm. (B) Relative fluorescent intensity of hnRNP F. The total green fluorescence of hnRNP F in the cells as semi-quantified by ImageJ software. n=3. ns *p* > 0.05. (C) The ratio of nuclear hnRNP F / the cytosol hnRNP F. Nucleic hnRNP F fluorescence intensity and the cytosol hnRNP F were separately semi-quantified by ImageJ software. The ratio of [nucleic ImRNP F]/ [cytosol linRNP F] was calculated. n=3, \**p* < 0.05. (D) HnRNP F expression measured by Western blotting. The expression of hnRNP F from transfected DCs was detected by western blotting. The relative expression of linRNP F were semi-quantified by measuring the density of western blotting bands. β -actin was used as a loading control. The ratio of hnRNP F/actin was calculated and normalized with NC controls. n=5. ns: *p* > 0.05.

## Discussion

Since their discovery in 1973 more subsets of DCs have been identified and more roles of DCs in disease development and progress have been recognized(23, 24). Knowledge about DC development and differentiation has accumulated(25-27). In this study, we demonstrated that circRNA is involved in the development of immune reactive DCs, expanding on our previous study(20). We, for the first time, showed that exonic circAEBP2 is expressed in the cytoplasm of DCs cultured from BM progenitors. The circAEBP2s levels increased in *in vitro*-cultured DCs over time and DCs cultured for 8 days and presenting a mature DC phenotype expressed much greater circAEBP2 than imDCs. LPS, a maturation factor, promoted DCs to become mature and upregulated the level of circAEBP2 measured by qRT-PCR, which is aligned with our preliminary microarray results (unpublished). The expression level of circAEBP2 was positively correlated with maturation status of DCs.

imDCs developed from BM progenitor cells are often characterized by low levels of MHC molecules, costimulatory molecules, and proinflammatory cytokines(28). Under stimulation, imDCs evolve into mDCs with high MHC I/II expression and costimulatory molecules CD40, CD80, CD83, and CD86(29). There are many signaling pathways and molecules participating in DC status transition(14, 30). In this study, we designed two different siRNAs (#1 and #2) targeting the conjunction sequence of circAEBP RNA and both of them knocked down of circAEBP2, however, siRNA #1, not siRNA#2 reduced linear AEBP2. This result shows that it is feasible to specifically and exclusively knock down circAEBP2. Therefore, we chose siRNA #2 to investigate the effect of circAEBP2 on DC development and function. We found that knockdown of circAEBP2 reduced the level of MHC II, CD40, CD80, and CD83 in DCs, indicating that knockdown of circAEBP2 arrests DCs at immature status and induces imDCs. Moreover, we found that the level of P-p65 was significantly reduced by circAEBP2 siRNA, indicating that the activation of the NFκB pathway (essential to DC maturation) is weakened by circAEBP2 siRNA. In contrast, inhibitory molecule CD200R1 expression was increased by circAEBP2 siRNA. Taken together, these data suggest that circAEBP2 is required for DC maturation and participates in DC development. Downregulation of circAEBP2 could induce imDCs. The current study, for the first time, demonstrates that circAEBP2 participates in the development of DCs. Manipulation of the expression of circAEBP2 could inhibit DC development and function.

The most important function of DCs is to activate naïve T cells. There are three essential signals for DCs to activate T cells(28). Signal 1 requires antigen presentation in the context of self-MHC for initiating recognition and priming naïve T cells. Signal 2 involves costimulatory molecules such as CD80 and CD86 and is essential for proper T cell activation. Signal 3 involves proinflammatory cytokines such as IL-12 that can further induce T cell proliferation (31). Lacking either one of these signals, DCs fail to robustly activate T cells. In this study, we determined the capacity of circAEBP2-silenced DCs to activate naïve T cells in an MLR *in vitro*. Our results show that knockdown of circAEBP2 impaired DC ability to activate naïve allogeneic T cells as evidenced by a reduction in T cell proliferation measured by a CSFE dilution assay, implying the circAEBP2 silenced DCs are immunosuppressive, not immune reactive, in terms of immune function. In general, immunosuppressive DCs can promote T_reg_ generation and induce immune tolerance(28). Thus, those DCs are also immune tolerogenic DCs (Tol-DCs). Tol-DCs have been used to treat autoimmune diseases and prevent immune rejection in organ transplantation(20, 32). In the present study, we co-cultured circAEBP2-silenced DCs with allogeneic naïve T cells *in vitro* and then measured inducible CD4^+^CD25^+^Foxp3^+^ Tregs. Our results showed that knockdown of circAEBP2 promoted DCs to generate more T_regs_. Taken together, our results suggest that knockdown of circAEBP2 can induce Tol-DCs, demonstrating that circRNA is a novel regulator for DC functions providing a novel strategy for *in vitro* generation of Tol-DCs. The potential of circAEBP2-silenced Tol-DCs in treating immune disordered diseases will be investigated in future, which is a limitation of the current study.

CircRNA functions through sponging miRNA and interaction with protein(33-36). To understand the molecular mechanisms by which circAEBP2 modulates DCs, we performed a series of RIP assays and found that circAEBP2 binds to cytosol hnRNP F. HnRNP F belongs to the hnRNP family(37) which is well known for its function in gene expression and signal transduction, including pre-mRNA and mature mRNA transcription, alternative splicing and polyadenylation(38, 39). Generally, hnRNP F is found to be broadly expressed in most tissues or cancer cells in the nucleus as well as in the cytoplasm(40, 41). Although there are no reports on hnRNP F in DCs, the current study shows that hnRNP F is highly expressed in DCs and expressed in both the nucleus and cytoplasm of mouse BM-derived DCs, as determined by ICC using hnRNP F Abs. Western blotting with hnRNP F Abs showed that knockdown of circAEBP2 did not affect the total hnRNP F protein. Furthermore, ICC result showed that knockdown of circAEBP2 reduced the ratio of cytosol hnRNP F/nucleic hnRNP F, indicating circAEBP2 binding to hnRNP F prevents the nucleic translocation of hnRNP F.

HnRNP F is an RNA binding protein and participates in various steps of RNA metabolism. It preferentially binds to RNAs with a poly(G)-rich sequence(42). To predict whether a pre-mRNA is bound by hnRNP F, a G-score is generated by the QGRS Mapper (http://bioinformatics.ramapo.edu/QGRS/) which aims to search Quadruplex forming G-Rich Sequences (QGRS) in a RNA sequence(43). The higher G-score a pre-mRNA has, the higher the possibility that hnRNP F binds to it. Using the QGRS Mapper, we calculated the G-scores for those changed proteins picked by MS between circAEBP2 silenced DCs and control DCs. We found that CD200R1 pre-mRNA had a higher G-scores, indicating that hnRNP F likely binds to CD200R1 pre-mRNA. However, it needs to further investigation whether and how circAEBP2 alters CD200R1 via hnRNPF. In the current study, we found that the level of CD200R1 protein was highly expressed in circAEBP2-silenced DCs in which nucleic hnRNP F was increased compared to NC siRNA-treated DCs. Our results also show that knockdown of circAEBP2 in DCs inactivated the NFκB pathway and impaired the capacity of DCs to activate allogeneic T cells. Therefore, we propose that circAEBP2 binds to cytosol hnRNP F, reducing the nucleic translocation of hnRNP F which binds to CD200R1, which in turn increases CD200R1 levels and leads to activation of the NFκB pathway and DC maturation.

In order to exclusively knockdown the circRNA while sparing its host gene, siRNA sequences have to cross the junction region of the circRNA, which impacts availability of siRNA sequences and robust silencing efficacy. This is a limitation of the study. Nevertheless, the current study clearly demonstrated the role of circAEBP2 in the development of DCs and the molecular mechanism by which circAEBP2 regulates DCs.

In summary, circAEBP2 participates in the development of DCs and modulates DC function through interacting with hnRNP F which regulates CD200R1-NFκB signaling, unveiling a novel regulator of DC development: circRNA. Knockdown of circAEBP2 induces Tol-DCs, providing a new strategy to generate Tol-DCs which can be used for controlling autoimmune diseases and immune rejection in organ transplantation. This study demonstrates an important role of circRNA in DCs and discover a new regulator for DCs, which increases our understanding on circRNA and DCs.

## Acknowledgments

The study was supported by grants provided by the Natural Sciences and Engineering Research Council of Canada (RGPIN-2019-04545) to XZ.

## Author contributions

QZ conducted experiments, analyzed data and wrote manuscript. BW and SL conducted experiments, WP and SMV provided scientific suggestions and participated in manuscript preparation. AG, participate manuscript preparation; XZ, formed research ideas and design and managed the study, manuscript writing and provided financial support.

