## Supplementary Tables for "Circular RNA AEBP2: A novel regulator in dendritic cell development"

**Supplementary Table 1. Primer sequences used for qRT-PCR in this study**

| **Gene/oligo** | **Forward Primer** | **Reverse Primer** |
| --- | --- | --- |
| CircAEBP2 | tgggtgaatgagagtgaacg | gagcgtccactggaaattgt |
| AEBP2 | agggaagggacacagtgttg | ctgcggcattgttctgtaaa |
| GAPDH | ggggtgaggccggtgctgagtat | cattggggtaggaacacggaagg |
| CircAEBP2 probe | gtgttgttttcc | ttatgagtact |
| Random probe | atgcaatggcacgtac | acggcatagatcgctc |
| CircAEBP2 probe template | gtgttgttttccacagtactcataagtgttgttttccacagtactcataa | |
| Random probe template | atgcaatggcacgtacgagcgatctatgccgaatgcaatggcacgtacgagcgatctatgccga | |

**Supplementary Table 2. Altered proteins in DCs transfected with circAEBP2 siRNA**

| Accession | circAEBP2 siRNA | NC Area | Fold change  (circAEBP2 siRNA v.s NC siRNA) |
| --- | --- | --- | --- |
| Q61164\|CTCF_MOUSE | 2.07E+05 | 8.38E+04 | 2.47 |
| Q8BYN3\|ITPK1_MOUSE | 8.75E+04 | 3.98E+04 | 2.20 |
| Q9ES57\|MO2R1_MOUSE | 9.09E+05 | 4.21E+05 | 2.16 |
| Q9D945\|CL031_MOUSE | 1.39E+06 | 6.72E+05 | 2.07 |
| P97930\|KTHY_MOUSE | 4.22E+05 | 8.32E+05 | 0.51 |
| Q9QVP9\|FAK2_MOUSE | 4.24E+05 | 8.44E+05 | 0.50 |
| Q99LJ0\|CT2NL_MOUSE | 1.96E+05 | 3.91E+05 | 0.50 |
| Q9QY24\|ZBP1_MOUSE | 5.52E+06 | 1.11E+07 | 0.50 |
| O09126\|SEM4D_MOUSE | 1.61E+05 | 3.24E+05 | 0.50 |
| Q9QUM4\|SLAF1_MOUSE | 2.99E+05 | 6.03E+05 | 0.50 |
| Q8CI95\|OSB11_MOUSE | 2.65E+05 | 5.42E+05 | 0.49 |
| Q91YN5\|UAP1_MOUSE | 7.37E+05 | 1.51E+06 | 0.49 |
| Q8R5F7\|IFIH1_MOUSE | 3.02E+06 | 6.27E+06 | 0.48 |
| Q9D0L8\|MCES_MOUSE | 8.31E+05 | 1.76E+06 | 0.47 |
| B1AZP2\|DLGP4_MOUSE | 1.18E+05 | 2.50E+05 | 0.47 |
| Q9DB43\|ZFPL1_MOUSE | 4.09E+05 | 8.76E+05 | 0.47 |
| Q9JHQ5\|LZTL1_MOUSE | 7.01E+04 | 1.51E+05 | 0.46 |
| Q3TTA7\|CBLB_MOUSE | 9.64E+04 | 2.08E+05 | 0.46 |
| Q91WV0\|NC2B_MOUSE | 1.05E+06 | 2.27E+06 | 0.46 |
| Q6PAM1\|TXLNA_MOUSE | 6.36E+05 | 1.40E+06 | 0.45 |
| Q921S7\|RM37_MOUSE | 2.60E+05 | 5.75E+05 | 0.45 |
| Q9D1C8\|VPS28_MOUSE | 3.30E+05 | 7.36E+05 | 0.45 |
| Q8CBA2\|SLFN5_MOUSE | 1.89E+06 | 4.40E+06 | 0.43 |
| Q8CE50\|SNX30_MOUSE | 1.39E+05 | 3.32E+05 | 0.42 |
| Q80TM9\|NISCH_MOUSE | 2.99E+05 | 7.34E+05 | 0.41 |
| Q60591\|NFAC2_MOUSE | 1.48E+05 | 3.76E+05 | 0.39 |
| Q64339\|UCRP_MOUSE | 3.81E+07 | 9.75E+07 | 0.39 |
| Q9QZQ1\|AFAD_MOUSE | 3.49E+05 | 9.02E+05 | 0.39 |
| P97360\|ETV6_MOUSE | 4.11E+05 | 1.07E+06 | 0.38 |
| Q9ERG2\|STRN3_MOUSE | 1.37E+05 | 3.60E+05 | 0.38 |
| Q3U481\|CS028_MOUSE | 1.16E+05 | 3.08E+05 | 0.38 |
| Q9ERS5\|PKHA2_MOUSE | 7.63E+05 | 2.03E+06 | 0.38 |
| Q6P4S6\|QSK_MOUSE | 2.45E+05 | 6.60E+05 | 0.37 |
| Q64282\|IFIT1_MOUSE | 4.70E+06 | 1.28E+07 | 0.37 |
| Q99JF8\|PSIP1_MOUSE | 9.67E+04 | 2.79E+05 | 0.35 |
| Q6A065\|CE170_MOUSE | 8.34E+04 | 2.88E+05 | 0.29 |
| Q5F2E7\|NUFP2_MOUSE | 6.34E+04 | 2.22E+05 | 0.29 |
| Q9CPN8\|IF2B3_MOUSE | 7.18E+04 | 2.74E+05 | 0.26 |
| P50096\|IMDH1_MOUSE | 9.35E+04 | 3.82E+05 | 0.24 |
| Q9D6K9\|LASS5_MOUSE | 1.31E+05 | 6.61E+05 | 0.20 |
